# Differences in external loads of different pitch types in Chinese male college baseball players

**DOI:** 10.1101/2025.06.26.661877

**Authors:** Shenglei Qin, Dingmeng Ren, Zheng Li, Bo Zhang

## Abstract

**Background:** In baseball, monitoring player load solely through pitch count and innings overlooks the variability introduced by different pitch types. This study analyzed differences in external load, measured as Player Load, across various pitch types using a GPS-based wearable device. Additionally, the relationship between Player Load and ball velocity was examined.

**Methods:** Four male collegiate Division I baseball players from China (age: 18.5 ± 0.58 years, height: 181.00 ± 6.83 cm, weight: 78.00 ± 5.48 kg) participated in this study. External loads were collected for four pitch types, fastball, changeup, curveball, and slider, using a wearable sensor device that recorded six variables: Maximum Player Load (Max PL), Maximum Rotation (Max Rot), Pitching Hourly Velocity, and Percentage of Acceleration Change in the Three Axes (Up Load %, Side Load %, and Fwd Load %). A Kruskal-Wallis non-parametric test was used to assess differences across pitch types, while stepwise multiple regression analyzed the impact of Player Load on pitching speed.

**Results:** Significant differences were observed among pitch types for Max PL, Max Rot, Hourly Speed, Up Load %, and Fwd Load % (p<0.05), while Side Load % did not show a significant difference (p>0.05). Stepwise multiple regression indicated that pitching speed was influenced by Max PL, with the equation: mph = 78.816 + 7.001 Max PL, R2 = 0.192, suggesting that Max PL accounted for 19.2% of the variability in pitching speed.

**Discussion:** External training loads in pitching vary by pitch type, with variable-speed pitches generating higher peak external loads, whereas fastballs are associated with greater ball speeds.

## Introduction

Quantifying an athlete’s training load is essential for improving performance and managing fatigue[1-4]. In baseball, there is limited knowledge about the key factors that contribute to an accurate assessment of pitching training load [5, 6]. Variables such as pitch count, pitch type, and pitching mechanics may influence training load [5]. Pitch count or innings pitched is commonly used as a proxy for training load in pitchers, but it does not correlate well with performance prediction or provide a precise measure of actual workload[5, 6]. Furthermore, relying on pitch count alone has proven less effective in preventing overuse injuries[7, 8]. Another limitation is that pitch counts fail to account for velocity variations between pitch types, which can lead to different levels of joint loading stress[9, 10]. This indicates that each pitch type may impose distinct demands on the body, and relying solely on pitch count may not accurately capture the cumulative workload.

The contribution of each pitch type to a pitcher’s overall training load may be reflected in the external training load associated with each pitch[11]. External training load can be quantified by analyzing the series of movements an athlete performs during training or competition[12]. This includes parameters such as distance covered, velocity, acceleration, deceleration, and impulse [13-15], among other factors. These movements can be further categorized into peak and cumulative external training loads [16]. Peak external training load is defined as the instantaneous measurement of these bodily movements[14], whereas cumulative external training load represents the total sum of an athlete’s movements over a training session or competition[17]. Global Positioning System (GPS)-based monitoring systems provide a means to quantify external training load using a metric known as Player Load[2]. This metric is derived from changes in acceleration across three axes: vertical, anterior-posterior, and lateral [18]. Research has demonstrated that these systems exhibit high inter-device reliability and strong convergent validity [2, 14, 17]. Due to their reliability, validity, and portability, these devices have been widely adopted in field sports such as football, Australian rugby, and rugby to monitor external training loads during both training sessions and competitions[12, 15, 16].

Sensor-based biomechanical monitoring has become increasingly common in baseball [19], with a primary focus on analyzing the mechanical characteristics of the shoulder and elbow joints in relation to pitching performance, including ball velocity and accuracy, as well as injury risk[19, 20]. However, research on the application of GPS-based wearable technology in baseball remains limited. To date, only one study has reported higher peak and cumulative Player Load in fastballs compared to curveballs, sliders, and changeups [11], suggesting a potential relationship between increased Player Load and greater ball speed. Additionally, variations in pitch types may result in differing triaxial acceleration loading patterns due to differences in pitching mechanics. Analyzing the peak Player Load associated with various pitch types, along with differences in triaxial acceleration, may provide a deeper understanding of how different external loads impact pitchers. This information could enable coaches to more accurately evaluate overall pitching loads, which may help optimize performance while managing fatigue.

The aim of this study was to examine differences in peak Player Load and triaxial acceleration loads across different baseball pitch types in collegiate male pitchers. The study also sought to explore the relationship between Player Load and pitch velocity. It was hypothesized that significant differences would be observed in peak Player Load and triaxial acceleration loads across pitch types, and that higher peak Player Load would be associated with increased pitch velocity.

## Materials and Methods

### Participants

Four collegiate baseball players (age: 18.5 ± 0.58 years, height: 181.00 ± 6.83 cm, weight: 78.00 ± 5.48 kg) participated in this study. All athletes were national-level competitors, right-handed pitchers, and had no major injuries or illnesses within the past six months. They were fully informed about the experimental procedures and voluntarily agreed to participate. The experimental protocol was approved by the regional ethics committee of Beijing Sport University (No. 2025202H). All participants provided written informed consent prior to the start of the experiment.

### Experimental equipment

Data collection was conducted using a GPS-based wearable device, the Catapult Vector S7 (Catapult Innovations, Melbourne, Australia), with each athlete assigned a unique module to prevent data overlap. Before each test, all subjects wore upper-body garments, with microsensors positioned between the scapulae. These microsensors recorded data at 100 Hz using IMU devices (*Figure 1*). External training load was quantified through a three-dimensional accelerometer-based formula, known as accumulated Player Load (PL), as provided by the manufacturer. This formula determines PL by calculating the square root of the sum of the squares of the instantaneous rate of change in acceleration across three orthogonal planes, anterior-posterior, medial-lateral, and vertical, divided by a scaling factor of 100, with results reported in arbitrary units (AU). The calculation formula is as follows. Given the frequent and rapid changes in activity and direction characteristic of basketball, existing research suggests that Player Load effectively measures the instantaneous rate of change in acceleration across three planes of motion[1].

Player Load is used to quantify the load experienced by a pitcher on each pitch. It represents a synthetic vector of the instantaneous rate of change in triaxial acceleration across different directions, divided by a scaling factor[18]. The calculation follows the formula:

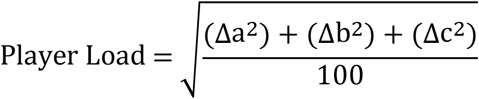

where Δa represents the instantaneous change in front-back acceleration (Fwd Load), Δb corresponds to the instantaneous change in medial-lateral acceleration (Side Load), and Δc denotes the instantaneous change in vertical acceleration (Up Load) [18]. Pitching speed (km/h) was measured using a radar gun (Stalker Sport 2, USA). The test metrics analyzed in this study included maximum Player Load (Max PL), maximum body rotation (Max Rot), percentage of triaxial acceleration change in different directions (Fwd Load%, Side Load%, and Up Load%), and pitching speed.

### Experimental design

Before the experiment, the GPS module was activated and securely fastened to the middle of the scapula using a vest. Each athlete completed a structured dynamic warm-up, which included jogging and stretching exercises. Following the warm-up, athletes were given the opportunity to throw an unlimited number of pitches across four pitch types, fastball, slider, changeup, and curveball, to ensure they were fully prepared for testing.

During the experiment, each pitch was thrown from the mound to a catcher positioned 18.44 m (60.5 ft) away. Pitchers were instructed to throw each pitch with maximum effort while the timing of each pitch was recorded. Pitch velocity was measured using a radar gun operated by surveyors positioned behind the catcher. There were no restrictions on pitch selection; before each pitch, pitchers verbally indicated the type of pitch they intended to throw. Each athlete completed 20 pitches per pitch type, resulting in a total of 320 recorded pitches from four athletes across four pitch types.

### Data Analysis

Descriptive statistics were presented as mean ± standard deviation (Mean ± SD). The variability of pitch type across six parameters, Max PL, Max Rot, Hourly Speed, Up Load%, Side Load%, and Fwd Load%, was analyzed using the Kruskal-Wallis non-parametric test. Stepwise multiple regression was used to assess the impact of pitching speed, with statistical significance set at *p* < 0.05. All statistical analyses were conducted using SPSS 25 (IBM SPSS Statistics, USA).

## Results

*Figure 2* illustrates the relationship between maximum Player Load (Max PL) and pitch speed across four baseball pitch types: changeup, curveball, slider, and line drive. The horizontal axis represents Max PL, while the vertical axis denotes pitch velocity (km/h). Each data point corresponds to a specific pitch type, differentiated by distinct colors and shapes. Fastballs exhibited the highest Max PL values, typically ranging from 4.00 to 5.00, and recorded the greatest pitch velocities, generally between 120.00 and 130.00 km/h. This suggests that fastballs require greater physical exertion to generate higher velocities. Sliders demonstrated slightly lower Max PL values, ranging between 3.50 and 4.50, with pitch speeds between 100.00 and

110.00 km/h, indicating that sliders are moderate-intensity pitches. Curveballs had even lower Max PL values, typically between 3.00 and 4.00, and produced the lowest pitch velocities among all pitch types, generally between 90.00 and 100.00 km/h. This indicates that curveballs impose a lower physical load on the body but result in correspondingly reduced speeds. Changeups recorded the highest Max PL values, usually ranging from 4.50 to 5.50, yet their pitch velocities were relatively low, typically between 80.00 and 90.00 km/h. Despite the greater physical load, the reduced pitch speeds suggest that changeups require higher exertion to create deception rather than velocity.

*Table 1* presents the descriptive statistics for all measured variables across the four pitch types.

**Table 1.** Descriptive statistics of variables under different pitching types (Mean ± SD)

|  | Fastball | Slide | Changeup | Curveball |
| --- | --- | --- | --- | --- |
| Max PL (AU) | 4.24±0.66 | 3.78±0.64 | 4.56±0.65 | 4.29±0.47 |
| Max Rot (AU) | 2.18±0.28 | 2.10±0.18 | 2.11±0.23 | 2.04±0.22 |
| Up Load% | 33.66±4.03 | 33.73±3.37 | 32.08±2.76 | 32.24±3.76 |
| Side Load% | 38.98±6.24 | 39.98±4.38 | 39.70±3.30 | 40.80±3.43 |
| Fwd Load% | 27.38±4.21 | 26.30±4.54 | 28.23±3.43 | 27.03±3.85 |
| Speed (km/h) | 121.53±4.68 | 100.06±6.70 | 113.14±2.88 | 98.60±4.59 |

Max PL: Max Player Load; Max Rotation

*Table 2* displays the differences in Side Load % across pitch types.

**Table 2.** Differences in external loads under different pitching types.

|  | Pitch Type Median M Quartile (P<sub>25</sub>, P<sub>75</sub>) |  |  |  | H value | p |
| --- | --- | --- | --- | --- | --- | --- |
|  | Change up(n=80) | Curveball (n=80) | Slide(n=80) | Fastball(n=80) |  |  |
| Max PL | 4.67 (4.2, 5.0) | 4.30 (4.0, 4.7) | 3.65 (3.3, 4.5) | 4.32 (3.6, 4.7) | 50.38 | 0.000** |
| Max Rot | 2.15 (1.9, 2.3) | 2.01 (1.8, 2.3) | 2.05 (2.0, 2.2) | 2.06 (2.0, 2.4) | 10.33 | 0.016* |
| Speed | 114.00<br>(113.0,116.0) | 100.00<br>(97.0, 101.0) | 100.00<br>(95.0, 105.8) | 122.00<br>(117.0, 126.0) | 254.68 | 0.000** |
| Up Load % | 33.00<br>(30.3, 34.0) | 32.00<br>(30.0, 35.0) | 34.00<br>(31.0, 36.8) | 33.50<br>(31.0, 37.0) | 14.47 | 0.002** |
| Side Load % | 40.00<br>(37.3, 42.0) | 41.00<br>(38.0, 44.0) | 40.00<br>(37.0, 43.0) | 39.50<br>(33.0, 45.0) | 4.60 | 0.204 |
| Fwd Load % | 29.00<br>(26.0, 31.0) | 28.00<br>(24.3, 30.0) | 26.00<br>(22.3, 29.0) | 28.00<br>(24.3, 31.0) | 11.58 | 0.009** |

\*p<0.05, \*\*p<0.01
Max PL: Max Player Load; Max Rot: Max Rotation

According to Table 2, no significant differences were observed in Side Load % among the different pitch types (*p* > 0.05). However, significant differences were found in Max PL, Max Rot, Hourly Speed, Up Load %, and Fwd Load % (*p* < 0.05), indicating that these parameters varied across pitch types. The specific median differences are detailed in Table 2.

The stepwise multiple regression fit equation is: mph = 78.816 + 7.001Max PL, R2 = 0.192, with Max Player Load explaining 19.2% of the variability in mph.

## Discussion

Traditional methods of assessing a baseball pitcher’s workload, such as pitch counts and innings pitched, have limitations in accurately reflecting total pitching load [6, 21]. Understanding the variations in external training loads across different pitch types may provide better insight into overall pitching demands. This study aimed to analyze differences in external load, specifically Player Load, and examine the relationship between peak Player Load and pitching velocity among collegiate male baseball players using GPS-based wearable devices. The findings revealed significant differences in Max PL, Max Rot, Hourly Velocity, Up Load %, and Fwd Load % across the four pitch types. Additionally, Max PL had a significant effect on Pitching Hourly Velocity, accounting for 19.2% of its variability. Due to its high velocity demands, the fastball requires a substantial physical load from the pitcher. This pitch type is typically employed to challenge batters quickly but necessitates a high level of strength and explosive power from the pitcher.

The slider falls between the fastball and curveball in terms of speed and physical load. It is a commonly used off-speed pitch that effectively disrupts batters’ timing. The curveball, on the other hand, requires less physical exertion, but its slower speed and curved trajectory make it a deceptive pitch for batters. Although the fastball generates higher maximum loads, its reduced velocity characteristics make it a strategic off-speed pitch that can deceive batters by altering velocity and movement.

Compared to previous research on external load differences among baseball pitch types, our findings reveal a broader range of variations. Earlier studies either found no significant differences in Max PL across pitch types [8] or reported that Max PL was only significantly higher for the fastball (4.0 ± 0.9) compared to the changeup (3.8 ± 0.9)[11]. In contrast, our study identified significant differences across all four pitch types. The highest Max PL was observed in the changeup (4.67; 4.2, 5.0), followed by the fastball (4.32; 3.6, 4.7), curveball (4.30; 4.0, 4.7), and slider (3.65; 3.3, 4.5). Additionally, the changeup exhibited high levels of Max Rot (2.15; 1.9, 2.3; *p* = 0.016). Significant differences (*p* < 0.05) were also observed in the percentage of acceleration along the longitudinal and anterior-posterior axes among the four pitch types. However, when examining pitch velocity, the fastball had a significantly higher hourly velocity (*p* < 0.001). Furthermore, Max PL had a significant positive effect on pitching velocity (R^2^ = 19.2), aligning with previous findings that higher pitch speeds are associated with increased external loading on the upper limb[16, 22, 23].

There is no consensus on the differences in external load, specifically Player Load, across various pitch types. One factor contributing to the variability in findings may be the placement of sensors [8, 11]. Previous studies have examined Player Load at three locations, the torso, upper arm, and forearm, across different pitch types, with significant differences observed only in the forearm. Variations in acceleration-based devices, sampling frequencies, and definitions of pitch onset and completion further complicate comparisons. Additional research is needed to better understand how external load differs by pitch type. Bullock et al. (2021) proposed using torso-mounted sensors as an alternative to pitch counts for identifying potential risk factors for pitching-related injuries [20]. However, this approach does not appear to be a reliable substitute for assessing loads on the shoulder and elbow [8]. In terms of trunk mechanics, all four pitch types are considered to follow similar movement patterns [11]. However, differences emerge at the forearm and wrist. Curveballs generate greater forearm rotation than sliders and fastballs [10], while both curveballs and sliders result in less wrist extension[10]. A systematic review by Grantham et al. (2015) found that curveballs produce lower proximal force and elbow torque compared to fastballs [24]. Consequently, using torso-mounted sensors may reduce the ability to detect mechanical load differences at the shoulder, elbow, and wrist across pitch types. Optical motion capture analyses have demonstrated that higher torso velocities are linked to reduced elbow torque[25-27]. The kinetic chain in the throwing motion functions by sequentially transferring energy along the body from proximal to distal segments, allowing the high output generated by the trunk to be converted into lower mechanical stress on the arm[8]. However, the relationship between trunk acceleration and mechanical stresses at the shoulder, elbow, and wrist remains unclear [8]. Wearable devices utilizing GPS, accelerometers, and gyroscopes are typically positioned on the back, approximately below the seventh cervical vertebra and between the scapulae. Because Player Load calculations primarily reflect movements of the body’s center of mass, where large fluctuations in instantaneous acceleration result in greater muscle contractions and energy expenditure[28], this metric is more commonly applied in whole-body running sports such as football and rugby [29]. In contrast, its applicability to unilateral kinetic chains, such as throwing, may be more limited.

Our study also suggests that Max PL plays a significant role in influencing pitch velocity. Nissen et al. (2009) reported that in youth pitchers, curveballs produced lower shoulder internal rotation moments and elbow valgus moments compared to fastballs, with these moments being influenced by both the magnitude of the applied force and the length of the force arm [30]. Among collegiate pitchers, an increase in maximum shoulder external rotation leads to greater rearward elevation during the arm-cocking phase, which increases stored elastic energy and muscle stretch. This energy is then utilized to maximize acceleration forces applied to the ball during the subsequent acceleration phase, ultimately increasing pitch velocity[31-33]. This transient interaction between force and the force arm may result in higher Player Load values in trunk movements, which in turn could contribute to greater pitch velocity.

This study was exploratory in nature and has certain research limitations. A larger sample size and multiple repetitions of pitch collection would have provided a more detailed assessment of external load differences among pitch types. Additionally, previous research has shown that factors such as body weight, leg length, and pitching stride length also influence pitching velocity [34]. Future studies should focus on larger sample sizes, including athletes of different competitive levels, while incorporating human morphology and biomechanical performance to increase the accuracy of analyses using torso-mounted sensors. Furthermore, investigating the effects of different pitch types and batters’ responses in actual game settings could provide insights that are more applicable to real-world baseball performance.

## Conclusion

This study demonstrated that external training loads in pitching vary based on pitch type. The findings revealed that variable-speed pitches generate greater peak external training loads, while fastballs result in higher ball velocities. Additionally, pitch velocity was influenced by the maximum Player Load at the time of release. Future research should consider sensor placement, human morphology, and biomechanical factors in studies involving larger sample sizes across different competitive levels. Further investigations should also examine peak and cumulative load variations between in-season and off-season training, as well as the physiological responses associated with pitching.

---

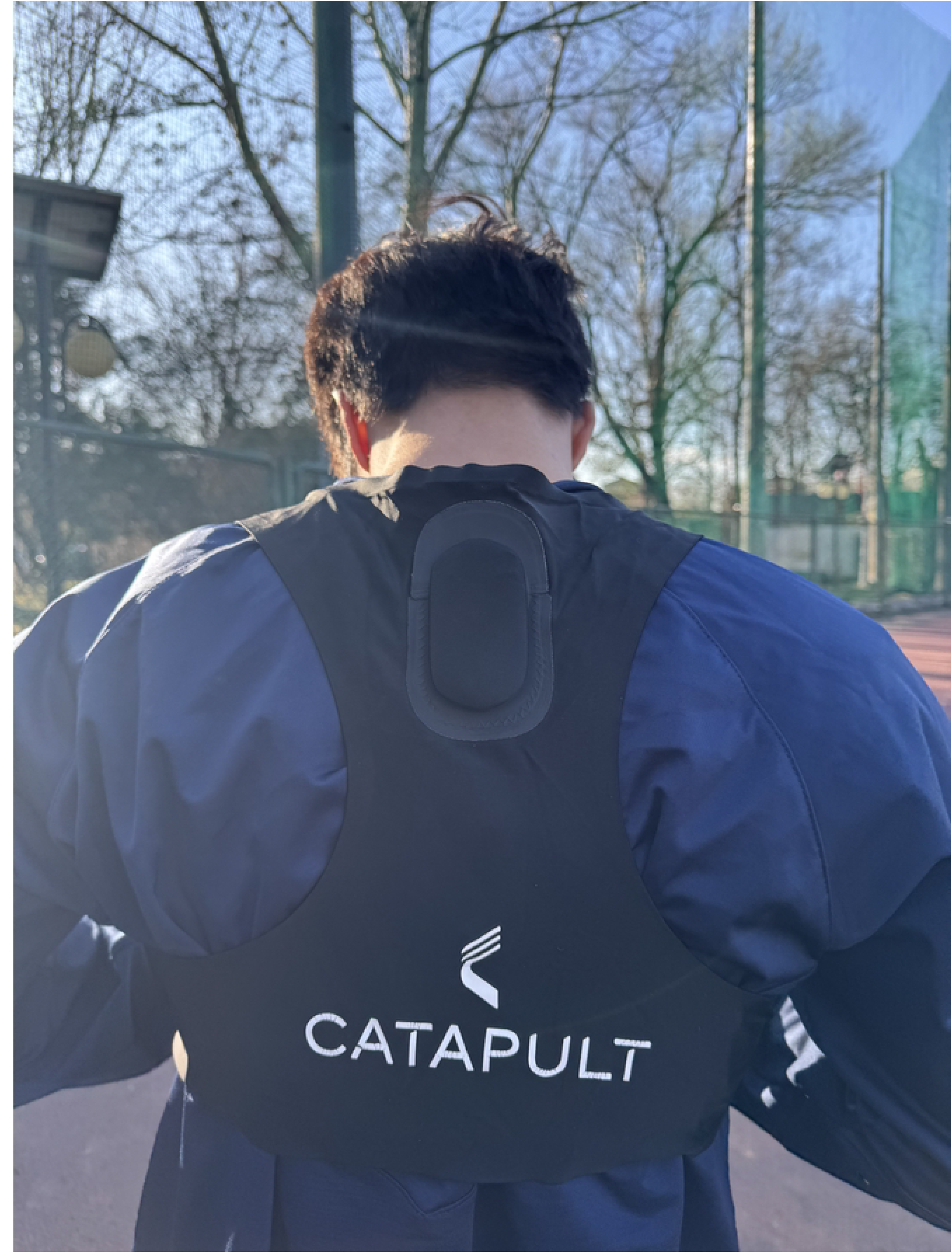

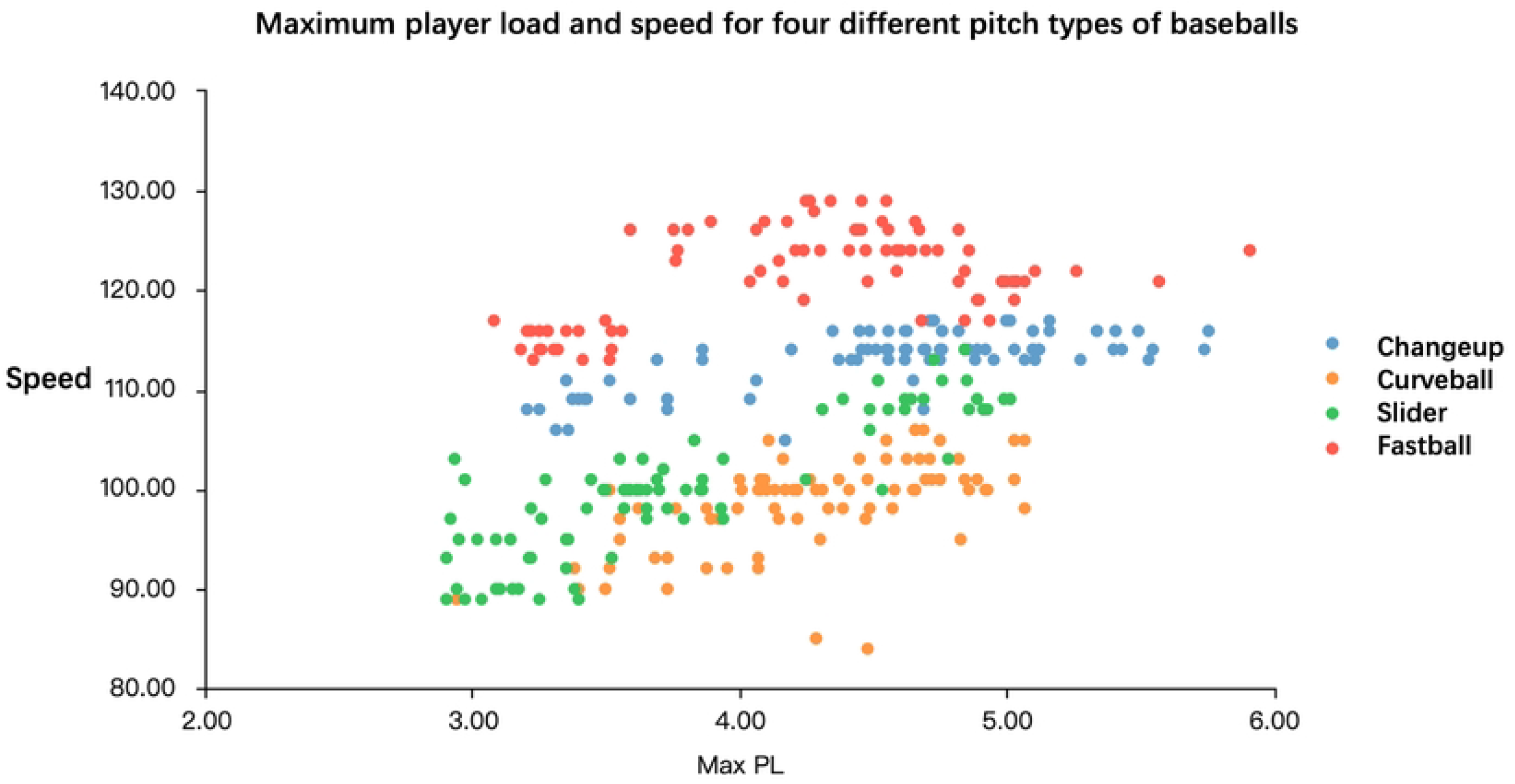

## Reference

1. Gabbet, T.J., QUANTIFYING THE PHYSICAL DEMANDS OF COLLISION SPORTS: DOES MICROSENSOR TECHNOLOGY MEASURE WHAT IT CLAIMS TO MEASURE? 2013.

2. Barrett, S., A. Midgley, and R. Lovell, PlayerLoad: reliability, convergent validity, and influence of unit position during treadmill running. Int J Sports Physiol Perform, 2014. 9(6): p. 945–52.

3. Bourdon, P.C., et al., Monitoring Athlete Training Loads: Consensus Statement. Int J Sports Physiol Perform, 2017. 12(Suppl 2): p. S2161–S2170.

4. Foster, C., J.A. Rodriguez-Marroyo, and J.J. de Koning, Monitoring Training Loads: The Past, the Present, and the Future. Int J Sports Physiol Perform, 2017. 12(Suppl 2): p. S22–S28.

5. Lyman, S., Effect of Pitch Type, Pitch Count, and Pitching Mechanics on Risk of Elbow and Shoulder Pain in Youth Baseball Pitchers. 2002.

6. Forman, J.C.B.A.S.L., THE IMPACT OF PITCH COUNTS AND DAYS OF REST ON PERFORMANCE AMONG MAJOR-LEAGUE BASEBALL PITCHERS. 2012.

7. Saltzman, B.M., et al., How many innings can we throw: does workload influence injury risk in Major League Baseball? An analysis of professional starting pitchers between 2010 and 2015. J Shoulder Elbow Surg, 2018. 27(8): p. 1386–1392.

8. Agresta, C., et al., Sensor Location Matters When Estimating Player Workload for Baseball Pitching. Sensors (Basel), 2022. 22(22).

9. Fleisig, G.S., et al., Kinetic comparison among the fastball, curveball, change-up, and slider in collegiate baseball pitchers. Am J Sports Med, 2006. 34(3): p. 423–30.

10. Fortenbaugh, D., G.S. Fleisig, and J.R. Andrews, Baseball pitching biomechanics in relation to injury risk and performance. Sports Health, 2009. 1(4): p. 314–20.

11. Bullock, G., et al., Comparative Pitching Biomechanics Among Adolescent Baseball Athletes: Are There Fundamental Differences Between Pitchers and Non-pitchers? Int J Sports Phys Ther, 2021. 16(2): p. 488–495.

12. Cummins, C., et al., Global positioning systems (GPS) and microtechnology sensors in team sports: a systematic review. Sports Med, 2013. 43(10): p. 1025–42.

13. Brendan R. Scott, R.G.L., Timothy J. Knight, Andrew C. Clark, A Comparison of Methods to Quantify the In-Season Training Load of Professional Soccer Players. 2013.

14. Varley, M.C., I.H. Fairweather, and R.J. Aughey, Validity and reliability of GPS for measuring instantaneous velocity during acceleration, deceleration, and constant motion. J Sports Sci, 2012. 30(2): p. 121–7.

15. Spencer, L.S.L.a.M., High Intensity Events in International Female Team Handball Matches. 2017.

16. Oyama, S., et al., Effect of excessive contralateral trunk tilt on pitching biomechanics and performance in high school baseball pitchers. Am J Sports Med, 2013. 41(10): p. 2430–8.

17. Johnston, R.J., VALIDITY AND INTERUNIT RELIABILITY OF 10 HZ AND 15 HZ GPS UNITS FOR ASSESSING ATHLETE MOVEMENT DEMANDS. 2014.

18. Luke J. Boyd, K.B., and Robert J. Aughey, The Reliability of MinimaxX Accelerometers for Measuring Physical Activity in Australian Football. 2011.

19. Yanagisawa, O., Alterations in pitching biomechanics and performance with an increasing number of pitches in baseball pitchers: A narrative review. PM R, 2024. 16(6): p. 632–643.

20. Bullock, G.S., et al., Differences in PlayerLoad and pitch type in collegiate baseball players. Sports Biomech, 2021. 20(8): p. 938–946.

21. Chalmers, P.N., et al., Fastball Pitch Velocity Helps Predict Ulnar Collateral Ligament Reconstruction in Major League Baseball Pitchers. Am J Sports Med, 2016. 44(8): p. 2130–5.

22. Glenn S. Fieisig, S.w.B., Rafael F. Escamilla and James R. Andrews, Biomechanics of Overhand Throwing with Implications for Injuries. 1996.

23. Hurd, W.J., et al., Pitch velocity is a predictor of medial elbow distraction forces in the uninjured high school-aged baseball pitcher. Sports Health, 2012. 4(5): p. 415–8.

24. Grantham, W.J., et al., The curveball as a risk factor for injury: a systematic review. Sports Health, 2015. 7(1): p. 19–26.

25. Arnel L. Aguinaldo, J.B., and Henry Chambers, Effects of Upper Trunk Rotation on Shoulder Joint Torque Among Baseball Pitchers of Various Levels. 2007.

26. Hirashima, M., et al., Kinetic chain of overarm throwing in terms of joint rotations revealed by induced acceleration analysis. J Biomech, 2008. 41(13): p. 2874–83.

27. Aguinaldo, A. and R. Escamilla, Segmental Power Analysis of Sequential Body Motion and Elbow Valgus Loading During Baseball Pitching: Comparison Between Professional and High School Baseball Players. Orthop J Sports Med, 2019. 7(2): p. 2325967119827924.

28. Wilson, R.P., et al., Moving towards acceleration for estimates of activity-specific metabolic rate in free-living animals: the case of the cormorant. J Anim Ecol, 2006. 75(5): p. 1081–90.

29. Dawson, L., Practitioner Usage, Applications, and Understanding of Wearable GPS and Accelerometer Technology in Team Sports. 2024.

30. Nissen, C.W., et al., A biomechanical comparison of the fastball and curveball in adolescent baseball pitchers. Am J Sports Med, 2009. 37(8): p. 1492–8.

31. Oyama, S., et al., Improper trunk rotation sequence is associated with increased maximal shoulder external rotation angle and shoulder joint force in high school baseball pitchers. Am J Sports Med, 2014. 42(9): p. 2089–94.

32. Tomoyuki Matsuo, R.F.E., Glenn S. Fleisig, Steven W. Barrentine, and James R. Andrews, Comparison of Kinematic and Temporal Parameters Between Different Pitch Velocity Groups. 2001.

33. Manzi, J.E., et al., The influence of shoulder abduction and external rotation on throwing arm kinetics in professional baseball pitchers. Shoulder Elbow, 2022. 14(1 Suppl): p. 90–98.

34. Manzi, J.E., et al., Kinematic Modeling of Pitch Velocity in High School and Professional Baseball Pitchers: Comparisons With the Literature. Orthop J Sports Med, 2024. 12(8): p. 23259671241262730.

